# An interplay of HSP-proteostasis, biomechanics and ECM-cell junctions ensures *C. elegans* astroglial architecture

**DOI:** 10.1101/2023.10.28.564505

**Authors:** Francesca Coraggio, Mahak Bhushan, Spyridon Roumeliotis, Francesca Caroti, Carlo Bevilacqua, Robert Prevedel, Georgia Rapti

**Author notes:** Authors contributed equally to this work (Mahak Bhushan, Spyridon Roumeliotis). Correspondence should be addressed to G.R.

## Abstract

Tissue integrity is sensitive to temperature, tension, age and is sustained throughout life by adaptive cell-autonomous or extrinsic mechanisms. Safeguarding the remarkably-complex architectures of neurons and glia ensures age-dependent, functional circuit integrity. Here we report mechanisms sustaining integrity of the *C. elegans* astrocyte-like CEPsh glia. We combine large-scale genetics with manipulation of genes, cells, and their environment, with quantitative imaging of cellular, subcellular features and material properties of tissues and extracellular matrix (ECM). We identify mutants with age-progressive, environment-dependent defects in glial architecture, consequent disruption of axons, synapses, and aging. Functional loss of epithelial Hsp70/Hsc70-cochaperone BAG2 causes ECM disruption, altered animal biomechanics, and hypersensitivity of glial cells to environmental temperature and mechanics. Glial-cell junctions ensure ECM-CEPsh glia-epithelia association. Modifying glial junctions or ECM mechanics safeguards glial integrity against disrupted BAG2-proteostasis. Overall, we present a finely-regulated interplay of proteostasis-ECM and cell junctions with conserved components that ensures age-progressively the robustness of glial architecture.

## Introduction

Maintaining lifelong tissue integrity across diverse environments is an outcome of the evolution of multicellular life. Interacting tissues must coordinate growth, shape changes, movement, and exerted forces in time and space. Cells are susceptible to temperature and metabolic changes, and continuously subject to mechanical loads, gravity, hydrostatic pressure, shear stress ^1–3^. Such factors influence animals at various scales, from molecular to tissue organization ^4–6^. To sustain tissue integrity, declining with age, cells need to adaptively respond to faced stresses by cell-autonomous or extrinsic mechanisms ^78^. They can respond to environmental factors and age progression by controlling composition of intercellular junctions or surrounding extracellular matrix (ECM) ^9,10^. Studies in epithelial tissues indicate that attachments to neighboring tissues or the ECM affect epithelial remodeling and force delivery to cells, proving critical for tissue organization ^5,6,11,12^. The complex mixture of ECM structural and functional proteins can provide spatio-temporal cues for sculpting epithelial morphogenesis and maintaining tissue organization ^5,13^. The effect of stressors on tissue architecture often depends on ECM composition and proteostasis ^4,10^, which can undergo remodeling through balanced production and turnover ^13,14^. Proteostasis needs to be adapted in age progression^10,19^ and declines in aging and neurodegenerative diseases ^7,15^, leading to cognitive dysfunction ^10,15,16^. ECM proteostasis can be regulated by temperature, metabolic shifts, and mechanical strains ^4,10^ and environmental features such as mechanics can affect epithelial tissue morphogenesis and integrity, throughout development ^5^. Investigating how proteostasis, ECM, and cell junctions integrate lifelong responses to external factors, to sustain tissue integrity against pathology, is key to comprehending tissue biology.

Nervous system tissues display a remarkable complexity of high-order cell interactions, and functions, that should be maintained throughout life. The interconnected circuit structure is primarily patterned during embryogenesis, through an interplay of cell recognition, glia-neuron interactions and molecular signaling, instructing precise positioning of neurons, glial cells, processes, and connections ^17–21^. Failure to preserve integrity of these highly specialized morphologies and connections results in compromised circuit functions, neuropathology and neurodegeneration ^1–3,22^. Therefore, it is crucial to understand the molecular mechanisms underlying maintenance of neuronal and glial cell architecture to ensure proper, lifelong circuit structure and function. Studies focusing on neurons’ susceptibility to mechanical stress, due to growth, movement, or injuries, identified mechanisms dedicated to neuron maintenance ^1,23,24^. In *C. elegans*, Ig-containing secreted or transmembrane proteins, L1 CAM and FGFR orthologs sustain positioning of neurons, axons ^24^, and synapses ^25^, while hemidesmosomes and spectrin protect axons from degeneration ^23^. Less is known about maintaining glial cell architecture. Specialized glia support neurons’ development, physical protection, repair, degeneration, and function ^26^. Similar to neuronal deficits, abnormalities in fine glial morphologies relate to neuropathology ^22^. Glial delicate, ramified processes formed in early development are vulnerable to mechanical strain and need to withstand elastic deformations during organismal growth. How glia face and withstand stress throughout life to sustain their architecture is poorly understood.

These questions can be addressed using *C. elegans*, a tractable model that allows large-scale genetics and single-gene manipulations. Its invariantly structured and mapped nervous system allows to study morphogenesis, maintenance, and interactions by visualizing and manipulating in single-cell resolution glia and neurons, and their subcellular features ^20,21,27^. Its brain neuropil, composed of 180 axons and 4 CEPsh glia, provides an *in vivo* platform to study circuit assembly and maintenance. We previously uncovered neuron-glia interactions underlying the assembly of functional circuit architecture. Embryonic CEPsh glia pioneer brain assembly, employing conserved cues to drive axon pathfinding of pioneer and follower neurons ^21^. Post-embryonic CEPsh glia also regulate synaptic positioning and neurotransmission ^28^. How CEPsh glia affect neuronal and axon integrity throughout life and growth, beyond synaptic distribution, is less clear.

In prior studies, we and others established that CEPsh glia and vertebrate astrocytes share functional, molecular, and morphological properties. They share transcription factors driving their fate ^28^, conserved guidance cues driving pathfinding ^21^, form tripartite synapses ^29^, influence synapse formation ^20^, regulate neurotransmission and present related transcriptomic profiles ^30^. On a larger scale, although not firmly sealing the neuropil, the CEPsh glial sheath provides a physical barrier between brain axons, basement membrane, adjacent tissues and the body cavity. This is reminiscent of blood-brain barriers in other species ^31,32^. CEPsh glia can provide a single-cell-resolution experimental system to study *in vivo* molecular mechanisms of astroglia architecture.

CEPsh glia form a specialized architecture to support their function. We previously uncovered that CEPsh form early, radial-glia-like processes, later transforming into astrocyte-like, ramified membranes, associating with axons and synapses ^21,33^. While CEPsh interior membranes appose axons, their external side contacts epithelia and a basement membrane separating them from muscle cells ^33^. Certain cues from epithelial cell and the basement membrane regulate their location/ shape ^34,35^. Yet, through which glial factors CEPsh glia connect to their neighbors and how they maintain their complex architecture throughout life and environmental changes remains elusive.

To investigate mechanisms maintaining CEPsh glia architecture and interactions, we performed genetic screens and isolated mutants presenting ectopic CEPsh glia membranes, with abnormal architecture. These glial changes are age-progressing, environment-dependent, and result in neuron, axon, and synapse mispositioning and degeneration. The causal alteration affects the HSP-co-chaperone UNC-23/BAG2, which acts from epithelia to sustain ECM architecture and integrity of CEPsh glia membranes. CEPsh glia associate with ECM and epithelial junctions and present cell-junction components. Partially disrupting cell-junction components in CEPsh glia in *unc-23* mutants disconnects them from their disrupted neighbors and safeguards their integrity, as does manipulating the animal’s exoskeleton and the mechanics of the tissue’s environment. Therefore, we uncover a key interplay of ECM proteostasis, glial-cell junctions and environmental factors of temperature and material properties, that ensures age-progressive CEPsh glia integrity.

## Results

### Post-embryonic CEPsh glia grow congruently with neighbouring tissues to envelop the brain

We and others established that CEPsh glia drive development and function of the *C. elegans* brain named *nerve ring* ^21,28,30,35^. They pioneer brain embryonic assembly and affect neurite development ^21,28^, synapse localization ^20,35^, and neurotransmission ^30^. CEPsh glia develop specialized morphologies to support their functions. Each one has an anterior process and a posterior membrane sheath associating with axons and synapses (Fig. 1, S1). Embryonically they grow thin non-branching processes ^21^, which then expand, throughout life, to generate ramified sheaths (Fig. 1A-B). CEPsh glia growth follows animal growth, in congruence with neighboring epithelia and mesoderm. CEPsh glial sheath membranes are juxtaposed to the external epidermis (Fig. 1C, Movie S1). Dorsal, ventral and lateral epidermal ridges extending from the cuticle, juxtapose the CEPsh glial sheaths directly without a basement membrane in-between ^33^. Outside these ridges, the CEPsh glial sheath neighbors four muscle quadrants (Fig. 1D, Movie S2), separated from them by an ECM basement membrane ^33^. Underneath the epithelial and mesodermal layers, the post-embryonic ramified CEPsh glia membranes ensheath the neuropil (Fig. 1E, Movie S3), associating with individual axons (Fig. 1F). They also extend processes ensheathing synaptic terminals as observed by trans-synaptic labeling using GFP reconstitution across synaptic partners (Fig. 1G-H) and electron microscopy ^33^. The precise morphology of CEPsh glia is key for glial-neuron interactions in brain architecture and function.

**Figure 1.**
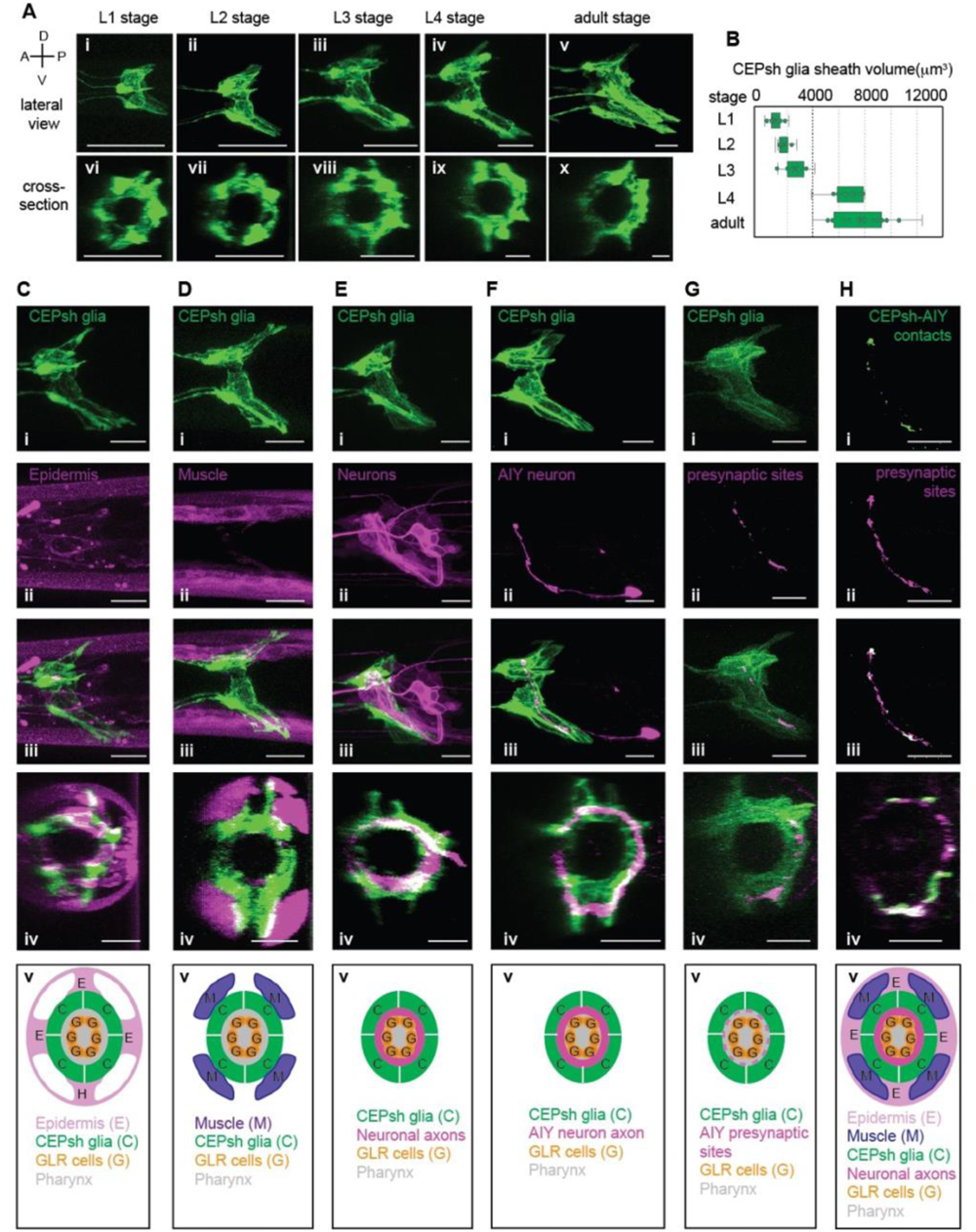
CEPsh glia grow congruently with muscle and epithelia, to envelop axons & synapses. **A-B.** Wild-type CEPsh glia grow throughout larval and adult stages, expanding their membrane volume. (B). n=14 animals per stage. **C-H.** CEPsh glia, in L4 animals, appose to epidermis (C), muscle (D), and envelop brain-neuropil axons (E), AIY axon (F) and RAB-3-labeled, AIY presynaptic sites (A-H) (CEPsh; green, others pseudocolored magenta). Cross-sectional views in C-Hiv, models in C-Hv. Reporters as listed in Methods, Tables S1, S3. Scale bars, 10μm. D, dorsal; V, ventral; A, anterior; P, posterior. (See Movies S1, S2, S3).

### CEPsh glia integrity suffers age-dependent disruption in *unc-23/BAG2* mutants

To uncover mechanisms regulating glial membrane architecture we performed a genetic screen, seeking mutants with abnormal CEPsh glial morphology (Methods). Among others, we isolated mutant *arg5*, exhibiting highly penetrant defects in CEPsh glial morphology with membranes occupying ectopic posterior territories up to 3 times farther from the cell body compared to developmentally synchronized wild-type animals (Fig. 2A-C). To identify the causal mutation, we subjected *arg5* mutants to whole genome sequencing, SNP mapping, and rescue experiments (Methods). We identified a G to A mutation in the *unc-23* gene, altering a conserved glutamic acid to a lysine in the dimerization domain of its vertebrate protein homolog ^36^. CEPsh glial defects of *arg5* are mimicked by the canonical allele *unc-23(e25)* ^41^and rescued when providing wild-type *unc-23* genomic sequences (Fig. 2D). Thus, proper CEPsh glial morphology requires UNC-23, the *C. elegans* homolog of vertebrate cochaperone BAG2 or Bcl-2-Associated Athanogene 2 that was previously implicated in regulating muscle architecture ^37,38^.

**Figure 2.**
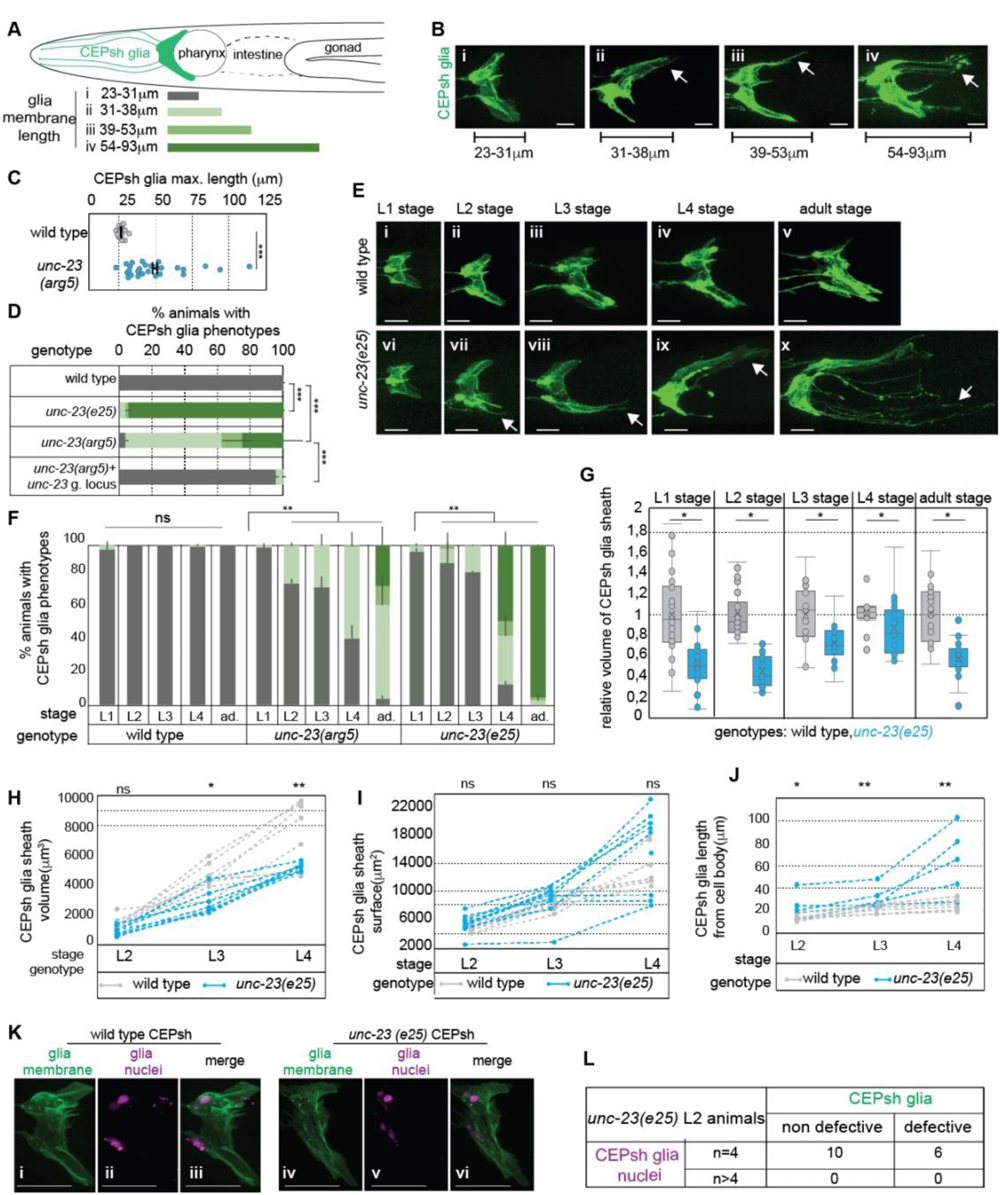
CEPsh glia present age-progressing loss of membrane sheath integrity in *unc-23* mutants. **A-C.** CEPsh glia (A, schematics), have defective morphology in *unc-23* L4 mutants (B), with increased membrane length (C). **D.** *arg5* defects are mimicked by *unc-23(e25)* mutant and rescued by the wild-type *unc-23* genomic locus (g. locus). **E-F.** Glia length defects aggravate with age progression. **D,F.** Error bars, mean ± standard deviation, n≥ 3 independent experiments, ≥ 100 animals total. ad, adults. **G-J.** Compared to wild-type animals, in *unc-23* mutants, CEPsh glia membranes throughout stages show abnormal decrease of volume (G,H) and increase of their length but not their surface, as assessed quantitatively (G, n= 20 animals per genotype) or in time-lapse experiments (H-J, n≥ 10 animals per genotype). **K-L.** CEPsh glia have normal nuclei numbers in L2 stage of *unc-23* mutant animals, at the start of establishment of their membrane defects. **B,E,K.** glia membrane, green; glia nuclei, magenta. Arrows, membrane defects. Arrowhead, nuclei defects. P-values: *** for ≤0,0001; **,≤0,001, *,≤0,01; unpaired test (C,G,I), Chi-square test (D-F). Scale bar, animal axes, reporters as in Fig. 1. (See Fig. S1, S2).

To understand the UNC-23/BAG2 function, we examined the timing and nature of the observed glial defects. Interestingly, CEPsh glia in *unc-23* mutants present age-dependent defects. In the 1^st^ larvae stage (L1), they have normal architecture compared to wild-type animals but show mild defects at the 2^nd^ larvae stage (L2) which increasingly aggravate in subsequent stages (Fig. 2E-F). To understand how defects arise, we examined if ectopic membranes result from abnormal pattern of cell divisions, or membrane growth or retraction, by quantifying cell numbers, membrane volume, surface, length, and growth rates in developmentally synchronized animals. CEPsh glia membranes in *unc-23* mutants compared to wild-type animals show extended length and decreased volume (Fig. 2D, G). To quantify glial growth rate we adapted immobilization protocols for long-term imaging and performed time-lapse imaging for individual animals throughout postembryonic stages (Methods, Movies S4-S5). Time-lapse quantifications revealed that CEPsh glia membranes grow continuously without retraction throughout larval stages and with comparable growth rate in wild-type animals and *unc-23* mutants (Fig. 2H). CEPsh glia membrane from L2 stage onwards have the same surface but increased length and decreased volume in *unc-23* mutants compared to wild-type animals (Fig. 2H-J). CEPsh glia present the same number of nuclei and anterior processes in *unc-23* mutants and wild-type L2 animals, before membrane defects are established. (Fig. 2K-L, S1), indicating that these embryonically born CEPsh glia do not suffer aberrant divisions or fate transformations. Thus, CEPsh glia in *unc-23* mutants stretch abnormally throughout animal’s growth. In L4 *unc-23* mutants, with largely established membrane defects, CEPsh glial cell nuclei are misaligned, mispositioned, and sometimes appear fragmented (Fig. 2M, S2). Overall, BAG2/UNC-23 is required for maintaining CEPsh glia integrity and its functional loss results in age-progressing loss of glial architecture with less compact, filamentous, discontinuous membranes, occupying ectopic territories (Movies S4-S5). The integrity defects of CEPsh glia in *unc-23* mutants are specific to axon-associated membranes since their anterior processes and tips have the same morphology in *unc-23* mutants and wild-type animals (Fig. S1).

### Age-progressing glia disruption results in abnormal circuit architecture and aging

CEPsh glia envelop axons and synapses throughout life and affect AIY neuron synapses ^20,21,35^. Thus, we assessed if age-progressive CEPsh glia integrity affects circuit architecture. The presynaptic sites of AIY interneurons and their CEPsh-glial contacts show decreased density and are sparser, occupying extended areas in *unc-23* mutants compared to wild-type animals (Fig. 3A-C). We queried whether these defects are specific to AIY synapses or relate to overall changes in circuit architecture. Importantly, increased axon length and cell-body mispositioning is observed in AIY and sensory neurons ensheathed by CEPsh glia in *unc-23* mutants compared to wild-type animals (Fig. 3D-H, N). CEPsh glia are implicated in age-dependent degeneration of dopaminergic neurons ^39^ Interestingly, *unc-23* mutants with age-dependent CEPsh glia disruption feature degenerating phenotypes of missing or shrunken cell bodies specifically in dopaminergic neurons CEP (Fig. 3J-L). The observed defects in sensory and CEP neurons correlate significantly with CEPsh glial defects (Fig. 3I). Importantly, circuit defects depend on disrupted CEPsh glia in *unc-23* mutants; amphid neuron defects are significantly suppressed when mutant CEPsh glia are ablated in L1, using an established system of caspase overexpression in glia (Fig. 3M, Methods). Finally, CEPsh glia were implicated in longevity ^40^. Aging *unc-23* mutant adults with defective CEPsh glia present decreased density of presynaptic vesicles (Fig. S3B-C), a hallmark of deteriorating neuronal healthspan ^41^, and decreased lifespan compared to wild-type animals (Fig. S3D). Thus, loss of CEPsh glia membrane integrity results in altered morphology of brain axons, concomitant synaptic defects and dopaminergic neurodegeneration. Altogether, our results highlight that age-dependent integrity of CEPsh glia is key to maintaining circuit architecture and healthspan, including features of axon positioning, synapse density, neurodegeneration, synapse aging, and animal lifespan.

**Figure 3.**
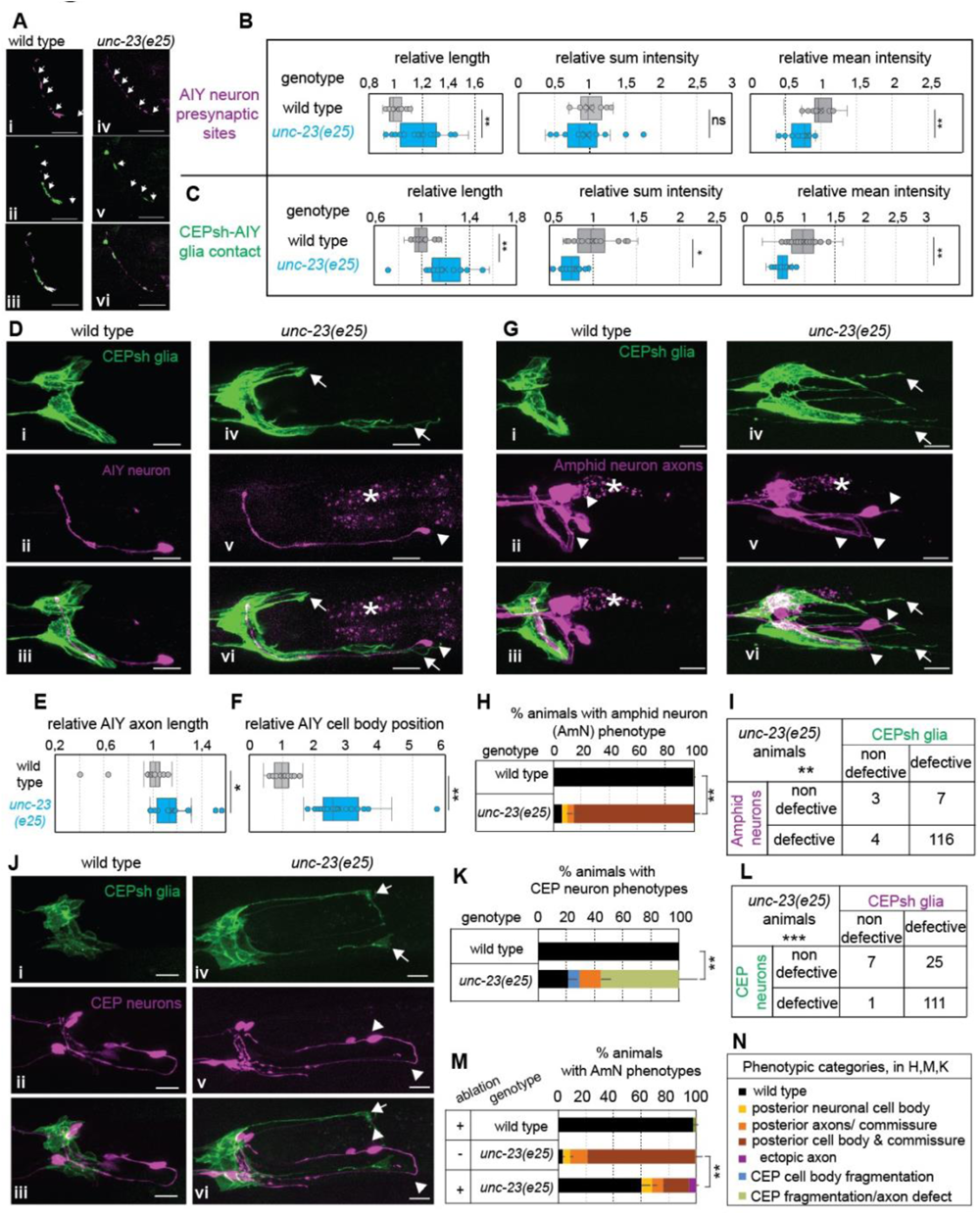
Synaptic and axon defects, and neurodegeneration are associated with abnormal glia integrity in *unc-23* mutants. **A-F.** Compared to wild-type animals, in *unc-23* mutants the RAB-3-labeled AIY presynaptic sites (magenta, A,B) and CEPsh-AIY contacts sites (green, A,C) are sparser, occupying larger areas with decreased density, while AIY (D-F) and amphid neurons (G-I) have defective positions and axon length, and CEP neurons show degeneration features (J-K) (CEPsh glia, green; all other pseudocolored in magenta, *, gut autofluorescence). **L-M.** Amphid neuron (AmN) axon defects (I) and CEP defects (L) in *unc-23* mutants correlate with glia defects. Axon defects in *unc-23* mutants are partially suppressed upon postembryonic CEPsh glia ablation (M). **N.** Panel explaining categories of defect types in panels H-M. **B, C, E, F.** n=20 animals per genotype. **H, K, M.** Error bars, mean ± standard deviation, n≥ 3 independent experiments, ≥ 120 animals total. P-values: ***, ≤0,0001; **, <0,001; *, <0,005; ns, non-significant. Chi-square test (H, K, M), unpaired t-test (B-C, E-F). Arrows, glia defects. Arrowhead, differences between mutant and wild-type in synapses (A), AIY (D), amphid (G), CEP (J). Scale bar, animal axes, reporters as in Fig. 1. Reporters as listed in Methods and Tables S3-5. (See Fig. S3).

### A conserved BAG2-Hsp70/Hsc70-DNAJB complex protects glial integrity in high temperatures

We set to examine the mechanism of action of UNC-23 in glial integrity. Its vertebrate homolog, cochaperone BAG2, interacts with stress-induced chaperone Hsp70 and its constitutively-expressed counterpart Hsc70, to regulate proteostasis ^42–44^. DNAJB cochaperones also regulate Hsp70/Hsc70, for substrate recognition, binding, and protein quality control ^43–45^. *C. elegans* DNJ-13/DNAJB1 and UNC-23/BAG2 can act oppositely for binding or release of HSP-1/Hsp70/Hsc70 substrates in other contexts ^37^. We assessed if such interactions of HSP-1/Hsp70/Hsc70, DNJ-13/DNAJB1 and UNC-23/BAG2 are involved in glial integrity. While *hsp-1* mutants show no defects in glia integrity, animals combining *unc-23* mutation with *hsp-1* mutation or knock-down of *hsp-1* or *dnj-13* restore glial integrity (Fig. 4B). Similarly, *hsp-1* and *dnj-13* mutations suppress muscle defects of *unc-23* mutants ^37^. Importantly, glial defects in *unc-23* mutant are rescued after expression of mouse BAG2 cDNA, suggesting a conserved molecular mechanism (Fig. 4C).

**Figure 4.**
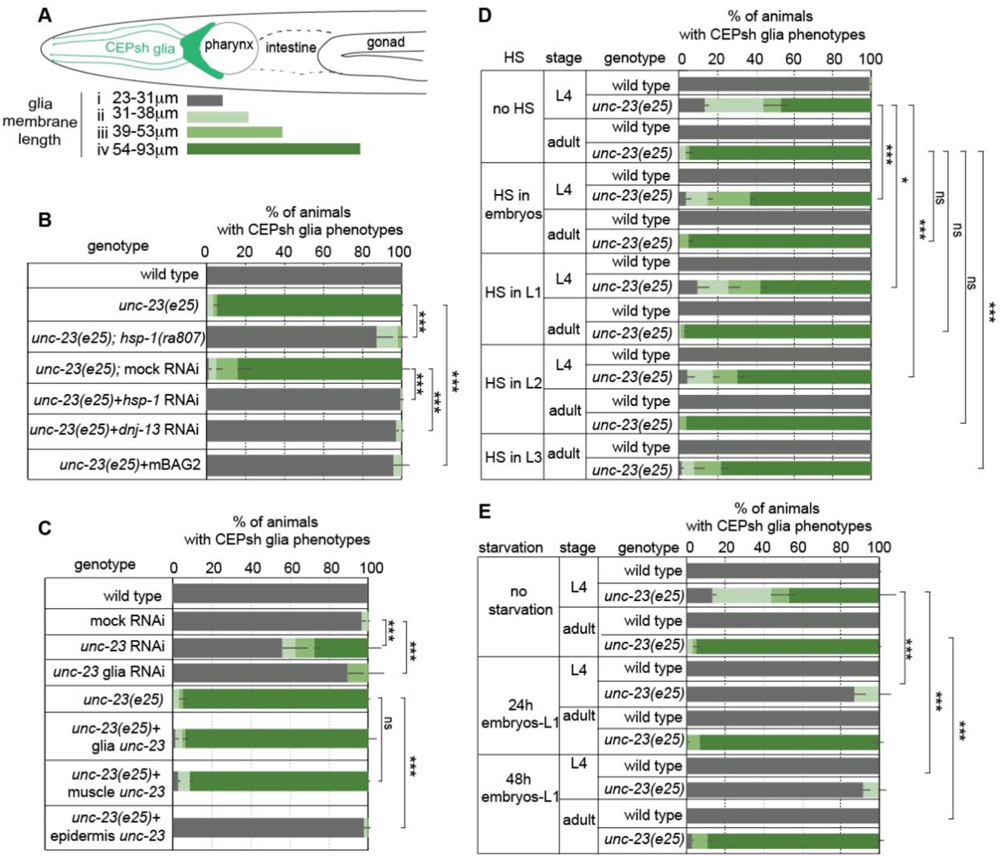
Epithelial UNC-23 acts with HSP-1 and DNJ-13 to regulate glial integrity in relation to temperature and nutrition. **A-B.** CEPsh glia defects (schematics, A) in *unc-23* mutants are suppressed by loss of HSP-1 or DNJ-13(B), rescued by mouse BAG2 (B), rescued by UNC-23 expression from epithelia, but not from glia or muscle (C), enhanced by temperature increase in embryos, L1, L2 and L3 (D), and partially suppressed by starvation (E). n≥ 3 independent experiments, ≥ 150 animals. HS, heat shock. Error bars, p-value, ns as in Fig. 2.

Hsp70/Hsc70 are implicated in temperature responses of cells ^46,47^, thus we assessed if thermal stress affects glia integrity. Interestingly, applying short-term heat-shock in *unc-23* mutants enhances the disruption of glial cell integrity (Methods, Fig. 4E). We also assessed if the effect of BAG2-Hsp70 in glial cell integrity is affected by nutritional states. Conversely to high temperature, starvation partially suppresses glial integrity phenotypes in *unc-23* mutants (Fig. 4F), suggesting that nutritional changes regulate UNC-23/BAG2 functions in this context. Overall, our results suggest that conserved functions of the UNC-23/BAG2-HSP-1/Hsp70/Hsc70 - DNJ-13/ DNAJB1 complex integrate tissue responses to temperature and nutritional changes to safeguard the integrity of glial cells and circuit architecture.

### Epithelial UNC-23/BAG2 protects glia integrity independently of epithelial integrity and allostery cues

To understand the site of action of UNC-23/BAG2 for glia integrity, we examined the presence of *unc-23* transcripts in transcriptomic data ^48^ and tested tissue-specific roles using RNA-interference (RNAi) and rescue experiments. *unc-23* transcripts in L2 larvae are mainly detected in epithelia, muscle, and glial cells including CEPsh. Bacteria-fed RNAi against *unc-23* in wild-type animals mimics *unc-23* mutant phenotypes (Fig. 4C). This suggests that UNC-23 does not act in neurons or glia, insensitive to bacteria-fed RNAi in wild-type animals. Moreover, vector-induced knock-down of *unc-23* in CEPsh glia does not induce glial defects (Fig. 4C). Specific expression of *unc-23* cDNA in epithelia, but not in CEPsh glia or muscle, and epithelial-expression of mouse BAG2 rescue the CEPsh glia defects in *unc-23* mutants (Fig. 4D). Thus, glia integrity requires conserved functions of UNC-23 as a BAG2 cochaperone in epithelia, that neighbor CEPsh glia.

Since UNC-23/BAG-2 acts in epithelia, we examined whether it is required for epithelial integrity or polarity, by visualizing membrane-bound fluorophores and the apical-surface SAX-7-derived marker ApiGreen ^27^. Importantly, the continuity of epithelial membrane and the intensity of apical-surface marker neighbouring the CEPsh glia is comparable in *unc-23* mutants and wild-type animals (Fig. S4A-B). Moreover, the integrity of epithelial cell junctions neighbouring the CEPsh glia appears similar in *unc-23* mutants and wild-type animals, as assessed by quantifying an enrichment of the apical-junction component DLG-1 (discs large MAGUK scaffold protein 1) neighbouring the CEPsh glia (Fig. S4C-E). Previous immunostaining studies proposed defects in epithelial hemidesmosomes in adult stages of *unc-23* mutants (Rahmani et al, 2015). While we cannot exclude *unc-23* mutant defects in other epithelial properties, domains or stages not assessed here, we detect no significant changes in epithelial polarity and cell junction integrity neighbouring the CEPsh glia that may account for impaired CEPsh glial integrity in *unc-23* mutants.

*C. elegans* epidermis regulates CEPsh glia through allostery cues CIMA-1/ SLC17A5 transporter and EGL-15/FGF receptor^35^. To investigate if UNC-23/BAG2 acts in the same pathway, we examined if changing levels of allostery cues affects the glial defects in *unc-23* mutants. Interestingly, neither overexpression nor genetic or RNAi-induced knock-down of CIMA-1/SCL17 or EGL-15/FGFR, suppresses the disrupted glial integrity of *unc-23* mutants (Fig. S5, Table S1). Therefore, UNC-23/BAG2 likely affects glial integrity independently of epithelial integrity and allostery cues, probably through new pathways of glia-epidermis communication.

### Ectopic glia sheath in *unc-23* mutants depends on ECM levels and accumulations

The BAG2-Hsp70/Hsc70 complex is implicated in proteostasis, protein folding, and disaggregation ^44,46^. Accordingly, the epithelial UNC-23/ BAG2, may safeguard glia integrity by regulating proteostasis required for epithelia-glia communication. We reasoned that if abnormal proteostasis of epithelial-secreted factors causes glial defects in *unc-23* mutants, modifying their levels may reverse these defects. To examine this hypothesis, we defined a predicted epithelial secretome, by the presence of signal peptide and epidermis-enriched transcripts in transcriptomes of L2 animals ^48^. We focused on transcripts of hyp7 epithelial cells, neighboring the CEPsh glia^49^. We performed an RNAi screen to knock-down epithelial-expressed ECM and secreted factors with epithelia-enriched transcripts (Methods, Table S1). We found that reducing the levels of ECM components UNC-52/Perlecan, EMB-9/COL4A5, and laminin LAM-1/LAMB1-2, partially suppresses the CEPsh glial defects of *unc-23* mutants (Fig. 5A). *unc-52* genetic mutation also suppresses CEPsh glial phenotypes in *unc-23* mutants.

**Figure 5.**
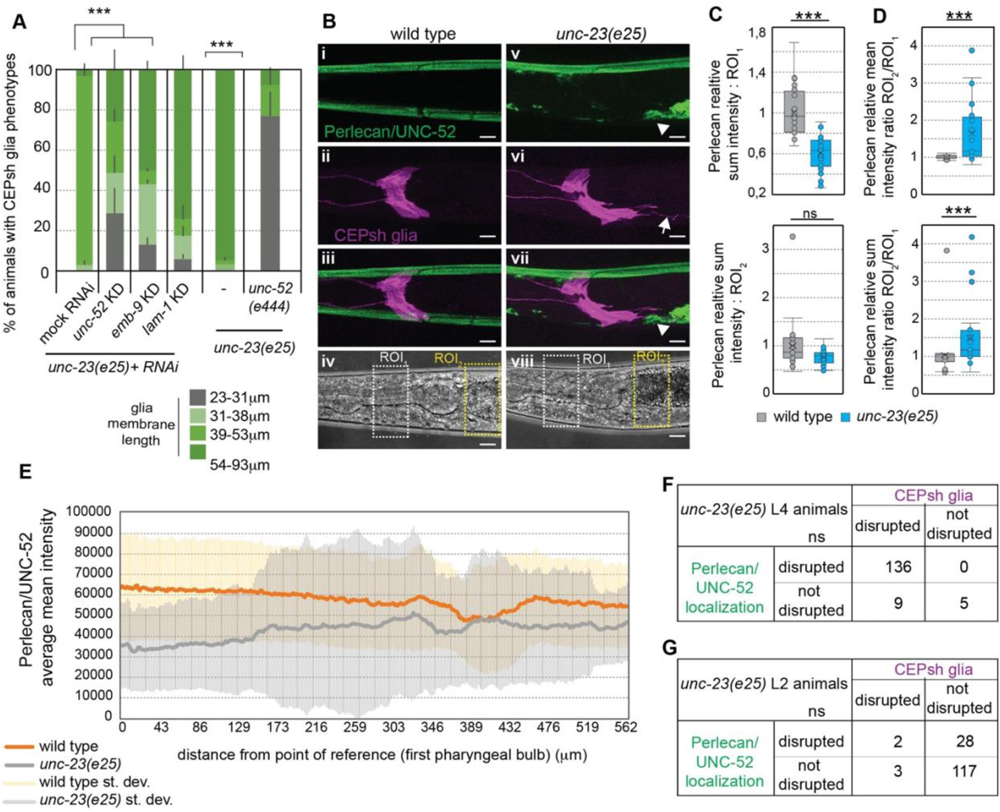
ECM is disrupted in *unc-23* mutants and altering its composition suppresses CEPsh glial defects. **A.** Knocking-down epithelial-expressed ECM components by RNAi (KD) or genetic mutant *unc-52(e444)* of the ECM component Perlecan partially suppresses CEPsh glia defects in *unc-23* mutants. n≥ 3 independent experiments, ≥150 animals total, Chi-square test. **B-E.** Perlecan/UNC-52 (green) has disrupted localization and accumulates posteriorly in *unc-23* mutants compared to wild-type L4 animals. n=20 animals per genotype (C-E). **E.** Perlecan intensity posterior to 1st pharyngeal bulb, represented by average (grey or orange) ± standard deviation (light grey or yellow) of 20 animals per genotype. **F-G.** Perlecan defects correlate to defects of CEPsh glia in L4 (**G**) but precede these defects in L2 animals (**F**), n ≥ 100 animals. Arrows, glia defects. Arrowhead, Perlecan defects. ROI_1_, ROI_2_ in B correspond to C-D. **(A-G).** Scale bars, animal axes, reporters, error bars, p-value, ns, as in Fig. 2. RNAi results in Table S1. (See Fig. S5).

To study ECM effects in glia integrity, we examined UNC-52/Perlecan and EMB-9/ COL4A5 using knock-in fluorescent tags ^50^. Compared to their continuous matrix in wild-type animals, the Perlecan and COL4A5 present abnormal localization in *unc-23* mutants, are depleted from areas neighboring normal CEPsh glia positions, and accumulate neighboring ectopic glial territories (Fig. 5B-E, S6). Importantly, Perlecan matrix abnormalities are significantly correlated with glial defects in *unc-23* mutant L4 larvae but precede these defects in L2 mutants (Fig. 5E-F). Overall, ECM components function downstream of UNC-23/BAG2 for glia integrity, they abnormally accumulate in *unc-23* mutants, correlated with and are followed by defective CEPsh glia, and altering their level suppresses CEPsh glial defects in *unc-23* mutants.

### CEPsh glia associate with epithelial junctions and ECM and glial-junction components affect glia integrity

Since ectopic CEPsh glia correlate with ECM accumulations, we reasoned that receptors of ECM may contribute to this association and the glial disrupion. To identify putative receptors of Perlecan, COL4A5, and Laminin, we queried the literature for interacting factors, presenting transcripts in CEPsh glia ^48,51^. Our queries highlighted proteins LET-805, DLG-1 and integrins, partaking in cell attachments. The fibronectin-domain transmembrane receptor LET-805, together with Matrilins MUA-3, MUP-4, compose hemidesmosomes while scaffolding protein DLG-1, and its binding partners AJM-1 and LET-413 compose adherens junctions, that mediate cell-ECM and cell-cell interactions respectively ^52^. These components are expressed in postembryonic CEPsh glia based on transcriptomics (Table S2). We examined if cell-junction components present specific localizations in CEPsh glia and may affect glial integrity. A fluorescently-tagged tool visualizing the DLG-1 binding partner AJM-1, expressed specifically in CEPsh glia, presents enrichments in the CEPsh glia membrane sheath in wild-type and *unc-23* mutant animals (Fig. 6A). This suggests that adherens junctions form in CEPsh glia membranes of normal and disrupted architecture (Fig. 6A). The CEPsh glia sheath also associates with DLG-1-containing epithelial adherens junctions, in wild-type and *unc-23* mutant animals (Fig. 6B-C). Glial-juxtaposed DLG-1 junctions associate closely with Perlecan in wild-type animals, and this association is maintained in *unc-23* mutants (Fig. 6D-E). Similarly to the CEPsh glia, epithelial DLG-1 junctions assume ectopic posterior localization in *unc-23* mutants, close to Perlecan accumulations (Fig. 6D-E).

**Figure 6.**
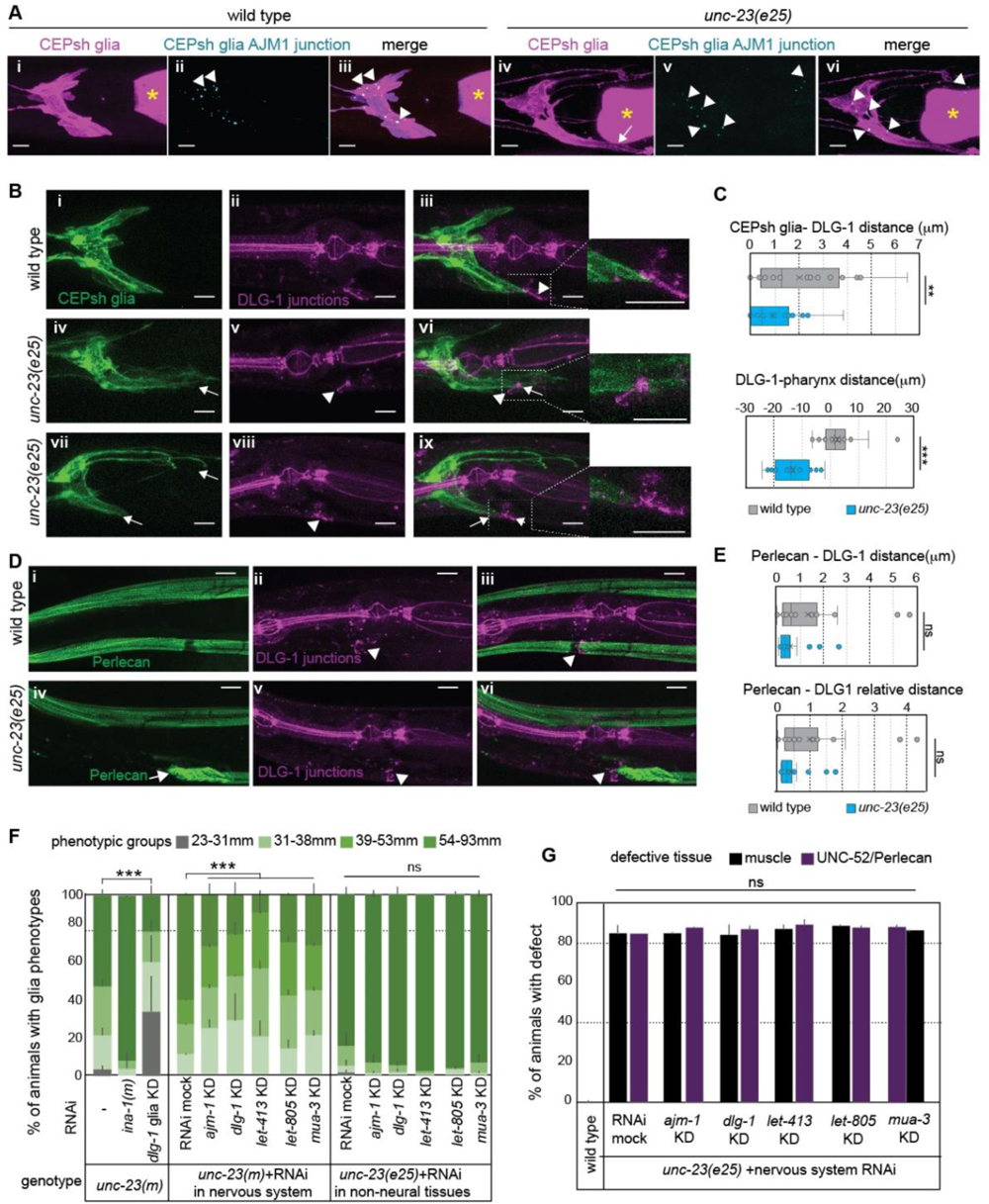
CEPsh glia appose to epithelial junctions and ECM, and disrupting glial junctions affects glia integrity in *unc-23* mutants. **A.** Adherens-junction marker AJM-1 (cyan) expressed in CEPsh glia (magenta) presents characteristic punctate localization in wild-type animals, largely maintained in *unc-23* mutants. **B-C.** CEPsh glia appose epithelial DLG-1 adherens-junction marker in wild-type animals and maintain their apposition, moving posteriorly in *unc-23* mutants. **D-E.** Epithelial DLG-1 closely associates with Perlecan matrix in *unc-23* mutants and wild-type animals. **F-G.** Vector-induced knock-down of DLG-1 specifically in CEPsh glia (glia KD) and bacteria-mediated RNAi knocking down of junctional components in CEPsh glia in the nervous system, but not in epithelia alone, partially suppresses CEPsh glia defects in *unc-23* mutants (F), while the Perlecan and muscle defects (G) are not recovered during this knock-down. Arrow, glia defects (A) or Perlecan/UNC-52 mislocalization (E). Arrowhead, DLG-1 enrichment (A, E). Yellow asterisk, pharyngeal bulb. **B-D-F.** n=20 animals per genotype, unpaired t-test. (G-H) n≥ 3 independent experiments, n≥ 100 total animals. *, Chi-square test. KD, knock-down. Scale bars, animal axes, reporters, error bars, p-values, ns, as in Fig. 2. RNAi results in Table S1.

We reasoned that if junctional components directly or indirectly connect the CEPsh glia to their neighbours (epithelia, ECM), knocking them down in *unc-23* mutants may release the glial membranes from these attachments, and suppress their presence in ectopic territories. To test this hypothesis, we knocked-down cell-junction components. Knocking-down DLG-1 specifically in postembryonic CEPsh glia, by vector-mediated RNAi, suppressed the ectopic CEPsh glia territories in *unc-23* mutants (Fig. 6F). Similarly, mutant CEPsh glia defects are partially suppressed upon bacteria-mediated knock-down of DLG-1 AJM-1, LET-413, LET-805, MUA-3 in the nervous system, using mutations that enhance RNAi-sensitivity in the nervous system (Fig. 6F, Methods). Transcripts of these components are largely absent from postembryonic neurons, thus this knock-down affects their glial transcripts. Conversely, CEPsh glia defects are not suppressed by knocking-down epithelial junctional components, using RNAi in *unc-23* mutants with RNAi-insensitive nervous system (Fig. 6F, Methods). This suggests that glia junctional components affect CEPsh glial integrity in *unc-23* mutants. Importantly, knocking-down these components in *unc-23* mutants does not suppress Perlecan and muscle integrity defects (Fig. 6G), thus it suppresses glia mislocalization not by recovering the ECM matrix but by disconnected it from the glia. Finally, mutating the integrin INA-1, presenting CEPsh glia transcripts, does not suppress CEPsh glia defects in *unc-23* mutants. In summary, CEPsh glia axon-enveloping membranes juxtapose epithelial adheres junctions closely associating with ECM and can harbour localized glial junction components, while interrupting these components releases the association of glia to abnormal ECM accumulations, safeguarding integrity of glial architecture away from ectopic territories.

### Material properties of the animal and its environment affect CEPsh glia integrity

To understand how glia and ECM deformations in *unc-23* mutants are affected by the environment, we altered the environment’s mechanical properties by changing the animals’ environment. Defective muscle integrity in *unc-23* mutants is suppressed when growing animals in liquid, but the underlying mechanism is unclear ^38^. CEPsh glia integrity is partially recovered upon growth of *unc-23* mutants in liquid. Interestingly, Perlecan matrix is also recovered, and its recovery is strongly correlated with recovery of glia architecture (Fig. 7A-B). Conversely, concurrent recovery of muscle integrity does not correlate with recovered glial integrity (Fig. 7C). Growth in liquid or on solid media presents differences in locomotory patterns and exerted forces. Animals adopt thrashing when swimming and crawling on solid media and exchange 1000-10,000 larger forces on solid compared to liquid media ^53,54^. To distinguish between effects of locomotion and forces, we induced paralysis on solid media to decrease the forces without causing thrashing. Such paralysis reverted the CEPsh glial defects of *unc-23* mutants (Fig. 7D). Thus, glial integrity upon BAG2/UNC-23 functional loss is sensitive to larger exerted forces.

**Figure 7.**
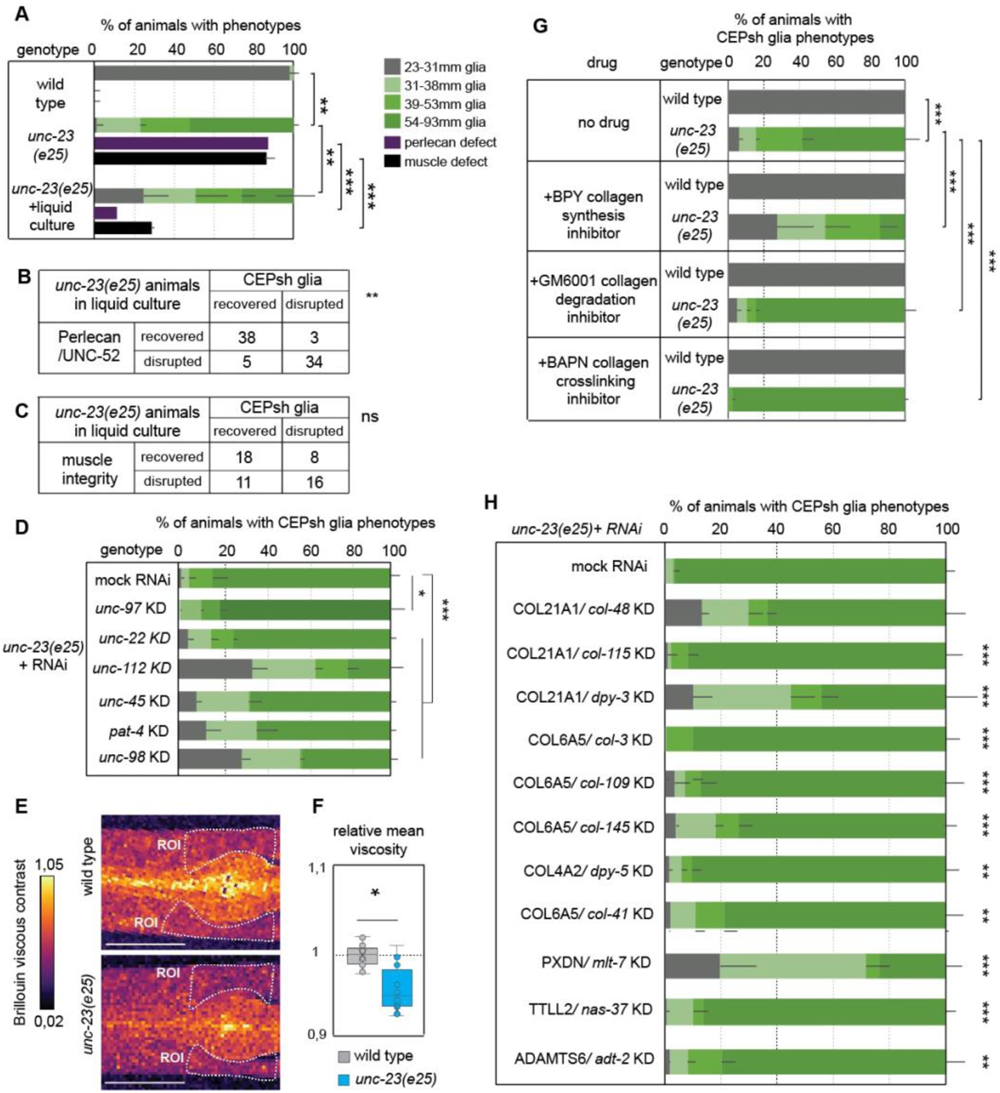
Altered animal’s and environment’s biomechanics, affect glia integrity in *unc-23* mutants. **A-C.** Growth of *unc-23* mutant animals in liquid media partially suppresses the defects of CEPsh glia, Perlecan matrix and muscle attachment. Liquid-induced recovery of CEPsh glia integrity correlates significantly with the recovery of Perlecan defects (B) but not with the recovery of muscle defects (C). **D.** Genetically-induced paralysis in *unc-23* mutants, by knocking-down (KD) genes of locomotory apparatus, partially suppresses CEPsh glia defects. **E-F.** L2 *unc-23* mutants show decreased tissue viscosity compared to wild-type. Doted area (D) is ROI (E). **F-G.** Collagen-modifying drug GM6001 and BAPN enhance CEPsh glia defects in *unc-23* mutants (G), while drug BPY or RNAi knock-down (KD) of collagens or collagen-modifying enzymes (H) partially suppress CEPsh glia defects in *unc-23* mutants. (A, D, G, H) n≥ 100 animals per genotype, chi-square test. (E) n ≥10 animals, unpaired t-test. Scale bars, animal axes, reporters, error bars, p-values, ns, as in Fig. 3. RNAi results in Table S1. (See Fig. S7).

In conjunction to exerted forces, tissue deformation depends on its material properties. All biological matter exhibits visco-elastic properties, displaying viscous and elastic characteristics when undergoing deformation. To measure *in vivo* biomechanical properties, we performed Brillouin microscopy, a non-destructive, label-free and contact-free method of optical elastography ^55^. Here, Brillouin microscopy measures both the so-called high-frequency elasticity as well as viscosity of biological samples, through the frequency shift and spectral linewidth of the Brillouin scattered photons interacting with the sample. We probed anesthetized *unc-23* mutant and wild-type animals, on L2 stage preceding the establishment of integrity defects. Interestingly, areas neighbouring CEPsh glia present decreased tissue viscosity in *unc-23* mutants compared to wild-type animals (Fig. 7E-F). Additionally, the animals’ tissue elasticity may present higher degree of variability in *unc-23* mutants compared to wild-type individuals (Fig. S7), as measured by the Brillouin elastic contrast ^56^. Thus, different animal biomechanics precede and underly age-dependent disruption of ECM and glia.

Since forces between tissues and their environment are transmitted through the animal’s exoskeleton, we reasoned that by modifying it we may regulate tissue integrity. To modify the cuticle, primarily composed by collagens, we used genetic manipulations of RNAi or chemical drugs BPY (collagen biosynthesis inhibitor), GM6001 (matrix metalloprotease inhibitor/collagen degradation inhibitor), and BAPN (collagen crosslinking inhibitor) (Methods). Interestingly, the disrupted CEPsh glia architecture in *unc-23* mutants is aggravated when chemically inhibiting collagen degradation or crosslinking (Fig. 7G, Table S1). Conversely, the disrupted CEPsh glia architecture in *unc-23* mutants is partially suppressed when inhibiting collagen biosynthesis chemically (using the above established drugs) or by knocking-down collagen-modifying enzymes ADT-2/ADAMTS, Peroxidasin-like MLT-7/PXDNL, pro-lysyl oxidase protease NAS-37 or specific collagens including DPY-3, DPY-5, COL-41, COL-43, COL-48, COL-109, COL-115, COL-145 (Fig. 7G-H, Table S1). Many of these genes have recognized predicted orthologues in mammals^57^. In conclusion, altering the material properties of the exoskeleton safeguards the age-dependent integrity of ECM and glia architecture.

Altogether, animals lacking epithelial Hsp70/Hsc70 cochaperone BAG2, suffer age-dependent disruption of ECM matrix, decreased tissue viscosity, and are hypersensitive to temperature, the environment’s material properties and tension forces. Their CEPsh glia follow mislocalized ECM matrix and epithelial junctions, through glial junctional-components, and experience abnormal stretching to occupy ectopic territories. Disordered glial integrity results in disruption of brain architecture after its establishment, with aberrant localization of neurons, axons, synapses and is associated with abnormal aging. Modifying glial junctions, the neighbouring ECM, material properties of the exoskeleton or the environment enhances the robustness of glia integrity against loss of BAG2.

## Discussion

Overall, this work uncovers a coordinated interplay between stress-dependent actions of the Hsp70-chaperone system, ECM architecture, cell-junctions and biomechanics, that regulates age-dependent integrity of astrocyte-like CEPsh glia and subsequent circuit architecture.

Upon functional loss of Hsp70-co-chaperone BAG2, the CEPsh glia suffer disrupted architecture with degeneration-like features, age-dependent shearing and nuclei fragmentation (Fig. 2), which is reminiscent of astrocyte gliodegeneration ^57^. CEPsh disrupted architecture affects axon-associated glial membranes (Fig. S1), and results in decreased neuron-glia contacts and density of presynaptic sites (Fig.3), also observed in mutants of allostery cues ^34^. We demonstrate that these synaptic defects are neither compartment-specific nor neuron-specific, but relate to global circuit defects. Disrupted CEPsh glial integrity causes neuron and axon mispositioning in diverse neurons, and dopaminergic neurodegeneration (Fig.3). In contrast, postembryonic killing of CEPsh glia does not cause defective axon morphologies ^30^. Thus, while absence of postembryonic CEPsh glia may not compromise circuit structure, their ectopic positioning disrupts its maintenance. These defects are succeeded by defective circuit aging (Fig.S3). Therefore, astroglia-like CEPsh glia are important for age-dependent, overall circuit maintenance. Similarly, ectopic activity of astrocytes, responding to injuries or aging, contributes to neuropathology ^22^. Gliodegeneration can precede aging and neurodegeneration and contributes to cognitive impairment ^7,57^. Uncovering mechanisms of astroglial integrity can shed light on understudied aspects of neuropathology.

Age-dependent disruption of CEPsh glia integrity arises from compromised BAG2-Hsp70/Hsc70 chaperone system (Fig.4). BAG proteins interacting with Hsp70/Hsc70 promote substrate release, to inhibit or enhance chaperone activity depending on the substrate and cellular context ^43^. The BAG2-HSP-system regulates CEPsh glia integrity from epithelia, by causing mislocalized ECM proteins and epithelial junctions that are followed ectopically by CEPsh glia (Fig. 4,5). Vertebrate BAG2 and Hsp70/Hsc70 homologs are implicated in neurodegeneration and aging ^9,10^, while in other contexts, they function in relation to temperature insults and nutrient changes ^12,58^. Their effects on cell integrity in response to environmental stress are less clear in the nervous system. Our findings establish that upon BAG2 loss, glia integrity is sensitive to heat and mechanical strain, while this is counteracted by nutritional block (Fig.4). This highlights that inefficient HSP-proteostasis affects the robustness of neural cell integrity upon temperature, nutrition, and biomechanic stress.

The BAG2-HSP complex acts for glia integrity, upstream of ECM and cell junctions. Upon BAG2 impairment, accumulations of basement membrane components (Fig.5) and mispositioning of epithelial junctions precede disrupted glial integrity and are followed ectopically by glia membranes (Fig.6). How CEPsh glia physically associate with epithelia and the ECM and how this affects their age-dependent integrity was previously unclear. Our work underscores that levels of Perlecan, Collagen IV, Laminin and glial cell-junction components are critical for association of glia to ECM and epithelial junctions (Fig.6). Manipulating these factors can discharge this association and safeguard glia architecture despite their neighbors’ mis-localization. Cytoplasmic cues supporting association to ECM/epithelia downstream of glial junctions, may include factors such as spectrins and intermediate filaments ^23^.

The intimate association of CEPsh glia with epithelia and basement membrane is reminiscent of interactions of astrocytes with endothelial cell junctions, or the Perlecan, Collagen IV and Laminin of the blood-brain-barrier (BBB) ^59^. These interactions are key for BBB integrity ^60^, and understanding which glial components sustain them is crucial. Our work implicates cell-junction components in an intimate glia-epithelia-ECM association, opening inroads to consider glial roles of their vertebrate homologs. Besides the suggested presence of desmosomes in astrocytes, in astrogliosis or Alzheimer’s, astrocyte functions of adherens junctions and hemidesmosomes remain unclear. Vertebrate DLG1 (Disc Large MAGUK Scaffold protein 1) is expressed in oligodendrocytes and endothelial cells and LET-413 homolog LRRC1 in astrocytes and oligodendrocytes (https://www.brainrnaseq.org/). Our work opens inroads to consider roles of these junctional factors in vertebrate glial cell architecture.

Epithelial-cell junctions affect integrity and robustness against internal and external forces ^6,11^, and facilitate tissue sliding during stretch ^6^. Stiffness and forces exerted on cells contribute to morphogenesis ^5^ while mechanical strain is implicated in neurodegeneration ^3^. Astrocytes are sensitive to mechanical tension ^61^, but how they integrate molecularly such experienced forces to sustain tissue integrity remains understudied. Our findings underscore how glial integrity relies on material properties. Impaired BAG2 is accompanied by reduced tissue viscosity and disruption of ECM components (Fig.6-7, S5) including of Collagen IV, a major contributor to ECM stiffness that affects local viscosity and the ability of tissues to withstand mechanical forces ^13^. Animals experience tension on the order of 10,000×G, 1000-10,000 times larger on high-viscosity compared to low-viscosity media while differences in undulations size and speed in these media are relatively modest. Following ECM disruption and decreased tissue viscosity, CEPsh glia integrity is more sensitive to high-viscosity than low-viscosity media (Fig.7), which is in line with effects of forces and biomechanics on other tissue deformations (Davidson., 2009).

*C. elegans* animals can be modeled as self-similar, shear-thinning objects, similarly to heart and brain, with gross mechanical properties governed by ECM components ^62^. Adding to internal ECM, specialized ECM composes the animal’s protective exoskeleton (cuticle) that contributes to force transmission ^63^. Physical properties of this ECM are determined by its collagen network ^62^, and its mechanics depends on its microstructure. Collagen fibers provide ECM and tissues with resistance to traction, and tensile strength to resist deformation, while the cuticle affects the animal’s bending stiffness ^62^. Collagen composition can affect biomechanics and shape changes upon applied forces. Here, reducing collagen levels partially safeguards age-dependent glial integrity, while blocking its degradation or crosslinking aggravates the glial defects in *unc-23* mutants (Fig.7). Thus, manipulating chemically or genetically the internal or external ECM surrounding the juxtaposed glia and epithelia, affects glial integrity regulated by HSP-proteostasis. Interestingly, our work highlights homologs of collagens or collagen-modifying enzymes associated accordingly with age-associated neuropathies ^64^ or ECM pathologies, glaucoma, glia and neurodevelopmental disorders ^65,66^.

Tissue responses to forces sometimes involve disaggregation or chaperoning of force-unfolded proteins and proteostasis may support robustness of ECM architecture upon mechanical stress ^10,67^. Meanwhile, *C. elegans* epithelia are involved in ECM deposition and force transmission^68^. Here, epithelial HSP-BAG2-proteostasis responds to mechanical stress upstream of ECM and cell junctions and its compromise causes age-dependent disruption of ECM and glial integrity, upon increased forces. Conversely, reducing the environment’s exerted forces, safeguards glia integrity despite impaired proteostasis (Fig.7). We uncover a proteostasis-tissue-biomechanics interplay that supports ECM and tissue robustness against mechanical strain. This interplay involves glial-cell junctions connecting glia to tissue and ECM neighbors, to maintain age-dependent integrity of their architecture and support circuit properties. Overall, our work paints a picture of well-balanced and tightly regulated protective mechanism that provides resilience of glial cells to environmental stressors.

## Materials & Methods

Further information and requests for information, reagent and resource sharing should be directed to and will be fulfilled by the Lead Contact, Georgia Rapti.

### C. elegans methods

*Animal Handling: C. elegans* strains are cultured as previously described ^21^. Bristol N2 strain is used as wild-type except when Hawaiian CB4856 is used for mapping. Unless otherwise indicated, animals are raised at 20°C, for two generations under *ad libitum* feeding conditions and L4 stage animals are scored. Animals used for phenotyping are hermaphrodites (not males).

*Germline Transformation:* Unstable extrachromosomal arrays are generated using standard microinjection protocols and integrations are performed using UV with psoralen (Sigma, T6137) and previously established protocols ^21^.

### Strains and plasmids used in this study

Previously published mutant alleles used in this study are provided by the CGC: LG.II : *unc-52(e444)*, LG.III : *ina-1(gm144)*, LG.IV : *hsp-1(ra807), cima-1(wy), eri-1(mg366)*, LG.V : *unc-23(e25)*, LG.X : *egl-15(n484), nre-1(hd20) lin-15B(hd126).* Information for newly generated alleles, mutants, extrachromosomal and integrated arrays is provided in Table S1. Information on unstable extrachromosomal and stably-integrated transgenes is provided in Tables S3-4 and used plasmid list in Table S4. Sequences of generated plasmids are available upon request.

### Isolation of mutants and genetic mapping

*Mutagenesis & Genetic Screening*: Animals are mutagenized using 70 mM ethyl methanesulfonate (EMS, Sigma) at 20 °C for 4 h. P0 generation of *nsIs374 LG.III* animals are mutagenized, F1-F2 generations are grown non-clonally. F2 progeny animals are examined on a epifluorescent microscope Zeiss Axioplan 2 (63×/1.4 NA objective, FITCH/GFP filter set (Chroma, Set 51019), or a fluorescent stereomicroscope Leica M165-FC (Objective Planapo 1.6x M-series and Filter set ET GFP -M205FA/M165FC). Animals with aberrant CEPsh glia morphologies are recovered.

*Gene Mapping:* Mutant *nsIs374* animals with ectopic CEPsh glia membranes (allele arg5) are crossed with wild-type males of CB4856 Hawaiian strain and recombinant progeny with CEPsh glia defects is isolated. Linkage mapping and SNP analysis highlighted linkage of mutant phenotype to a∼4.6-cM interval from LG V 0,46 cM (SNP K11C4.1: V: 6,898,091) to V 5,11 cM (SNP C26E1: V: 13,293,704). Whole-genome-sequencing detected mutations in 14 genes in this region. Mutants are rescued using fosmid WRM0626bA02, which includes *unc-23,* and tissue-specific expression of *unc-23* cDNA. Mutants are out-crossed at least four times.

### Animal synchronization

Animal synchronization for imaging experiments: Gravid adults are placed on OP50 plates, left for 2 hours to lay eggs, then removed. Laid progeny is grown at 20 °C until reaching L4 stage at 48 hours. L4 animals are imaged unless otherwise indicated.

Animal synchronization for RNAi/drug treatment experiments: Gravid adults are bleached, and recovered embryos are grown in M9 solution at 20C for at least 15h until L1 larval arrest. Synchronized larvae in L1 arrest are recovered and placed on plates with the relevant treatment.

### Heat-shock and starvation studies

Gravid adults are bleached, embryos are recovered and grown at 20 °C until desired stages are obtained. These synchronized animals are exposed to 30 °C for 5 hours during stage of interest. Controls are kept at 20 °C. After treatment, animals are grown at 20 °C to reach the stage of interest. For starvation studies, synchronized embryos are grown without bacteria and L1 animals are recovered on food, after 24h or 48h.

### Dye filling assays

Animals are collected from plates in M9 with 10 μg/mL of lipophilic dye 1,1′-dioctadecyl-3,3,3′,3′-tetramethylindocarbocyanine perchlorate (DiI, prepared in N,N-dimethylformamide), incubated in dye for 30 min in the dark, washed twice with M9, placed on bacteria plates for 3 hours. Amphid-commissure axon phenotypes are scored in epifluorescent microscope Zeiss Axiovert (63×/1.4 NA objective, Zeiss Filter set 43 HE Cy3). Different defects in amphid commissures are scored as described below.

### CEPsh glia ablation

Post-embryonic ablation of CEPsh glia is performed using stable genetic array (nsIs180) expressing a reconstituted Caspase-3 in CEPsh glia from the L1 larvae ^30^.

### Drug application

Plates containing drugs BPY, GM6001 and BAPN are prepared as follows: 2,2’-Bipyridine (BPY, Thermofisher Cat#030569) or GM6001 (Abcam, ab120845) is diluted in DMSO and 3-Amino-propionitril -fumarat (BAPN, Sigma, A3134) is diluted in H2O, each in final concentration 100mM. Drug-containing plates are prepared by adding drug in both the agar and bacteria, in final concentration of 100μM. Synchronized eggs are grown on drug plates, at 20 °C, until the L4 larval stage, then used for phenotypic analysis.

### Growth in liquid bacteria media

Embryos synchronized as above are grown at 20 °C in plates containing 1ml liquid bacteria culture and replenished with liquid media every 12 hours. Animals are scored phenotypically at L4 stage. Control animals are prepared identically but grown in solid culture.

### Studies of RNA interference

RNA interference (RNAi) is performed as previously described, using vectors expressing double-stranded RNA (dsRNA) or by feeding animals with dsRNA-expressing bacteria. In the first case, knocking-down transcripts in specific cell types is achieved by expressing dsRNA under cell-specific regulatory sequences. In the second case, knocking-down transcripts in nervous system cells is achieved by RNAi applied in genetic backgrounds combining mutations *eri-1(mg366)* and *nre-1(hd20) lin-15B(hd126)* that sensitize nervous system cells to RNAi. Absence of these genetic backgrounds, allow RNAi in other tissues (epithelia, muscle, intestine) without affecting nervous system cells. In this study, RNAi-bacteria-feeding is performed post-embryonically. Synchronized larvae after L1 arrest are fed *E. coli* HT115 with pL4440 vectors targeting specific genes, grown at 20 °C for 3 days and scored as adults unless otherwise indicated. RNAi clones from the Ahringer or Vidal libraries are used. Triplicates for each experiment are performed. RNAi clones presenting statistically significant effects were verified for specificity of the target using DNA sequencing. Imaging of animal phenotypes after RNAi applications is performed in Leica M165-FC (Objective Planapo 1.6x M-series and Filter sets ET GFP - M205FA/M165FC and ET RFP - M205FA/M165FC).

### *In vivo* imaging and phenotypic scoring/quantification

Live imaging is performed on postembryonic stages L1, L2, L3, L4. Animals are anesthetized using M9 buffer containing 20-25 mM sodium azide, mounted on pads of 2% agarose and image acquisition is performed immediately after. Imaging is performed with optical sections of 0.5 μm spacing for the following labeling:

CEPsh glia membranes or nuclei, epidermis, muscle, all axons, AIY neurons, CEP neurons are visualized *in vivo* using each of *Phlh-17::myrGFP* or *Phlh-17::myrGFP-SL2-NLS-mCherry, Pdpy-7::mKate, Pmyo-3::RFP*, *Prab-3::mKate-PH, Pttx-3::GFP, Pdat-1::GFP* respectively (Tables S4-5). Amphid neurons are visualized using dye filling. AIY presynaptic sites and CEPsh glia-AIY synapse contact sites are visualized using the markers *Pttx-3::mCherry::rab-3* and *Phlh-17::CD4::GFP(1-10) + Pttx-3::CD4::GFP(11)* (Tables S4-5) respectively. Length, area, intensity of fluorescent signals, and distances between fluorescent signals are quantified using Fiji software. Defects of number/position/fragmentation, axon position/ ectopic growth, of amphid or CEP neurons cell body are measured in population.

UNC-52/Perlecan and EMB-9/Collagen IV are visualized using each of CRISPR knock-in strain *unc-52(qy80[mNG+loxP (synthetic exon)::unc-52*]) or *emb-9 (qy24[emb-9::mNG+loxP])* respectively (Tables S3-4). Intensity of fluorescent signals in regions of interest is measured. Epithelial apical domains are visualized with ApiGreen-GFP marker *Phlh-17::SAX_7delcyt_sfGFP (*Tables S4-5), a construct derived from sequences of transmembrane protein SAX-7 localized exclusively to apical surfaces in a canonical epithelium ^27^. Region of interest for fluorescence signal is defined between the posterior end of the 1st pharyngeal bulb and the anterior of the 2nd pharyngeal bulb (based on images of the bright-field channel). Epithelial *DLG-1* is visualized using marker *Pdlg-1::dlg-1::*RFP (Tables S4-5), DLG-1 enrichment is quantified ventrally to the 2nd pharyngeal bulb. AJM-1 adherens junctions in CEPsh glia are visualized using marker *Phlh-17::ajm-1-cDNA-CFP* (Tables S4-5), via the CEPsh glia-specific expression of a previously established CFP-tagged label of AJM-1 protein localization domains ^27^. Animals are observed *in vivo* on a fluorescent stereomicroscope M165FC (Objective Planapo 1.6x M-series, Filter set and ET RFP – M205FA/M165FC) or an epifluorescent microscope Axiovert Zeiss (63×/1.4 NA objective, Zeiss Filter set 43 HE Cy3). Image acquisition is performed using a confocal microscope SP8 Leica (Objective and Filter as above) or a spinning disk Olympus iXplore SPIN SR (100×/1.35 NA objective, Filter set DM D405/488/561/640).

### Time-lapse imaging of postembryonic CEPsh glia

Post-embryonic live time-lapse imaging is performed on synchronized animals, immobilized using Polybeads Microspheres (Polysciences 00876-15), as described previously (^69^. No sodium azide is used. Images are acquired using spinning disk microscope Olympus iXplore SPIN SR. Immediately after imaging, animals are recovered by removal of coverslip, addition of M9 buffer, recovery on fresh bacteria plates and grown at 20 °C until the next stage. Individual animals are followed separately. Imaging and recovery are repeated for each of L2, L3, L4 stage.

### Brillouin microscopy and imaging of mechanical properties

Brillouin microscopy measured tissue mechanical properties such as elasticity and viscosity in the GHz frequency range through the interaction of light with the sample’s acoustic phonons ^55^. The shift and linewidth of the Brillouin scattered light spectrum gives information about the so-called longitudinal modulus which is directly related to the elastic and viscous modulus of the material, respectively. Brillouin imaging is performed using a Brillouin microscope previously described in^70^. Briefly, this consists of a commercial Zeiss body (Axiovert 200M) coupled with a home-built spectrometer based on a 2-VIPA configuration. A 532nm laser (Torus, Laser Quantum) is used for Brillouin imaging. An 488nm laser, coaligned with the 532nm laser, allows for confocal imaging of GFP fluorophores. Wild-type or *unc-23* mutant animals, synchronized at L2 stage, expressing markers of CEPsh glia or UNC-52::GFP are used. Animals are starved for 2 hours before experiments to minimize variability of Brillouin signal in the pharynx due to bacteria content. Animals are anesthetized using M9 buffer containing 5mM sodium azide and mounted on a slide of 2% agarose in 10mM sodium azide. Images of GFP are acquired before and after the Brillouin acquisition. Brillouin images are acquired with a 40x 1.0NA Zeiss objective and an integration time for a single point of 100ms. The optical power on the sample is kept below 4 mW and no apparent photodamage is observed after imaging.

### Image analysis and processing

#### For analysis of spinning disk/ confocal imaging

Length of distances between structures, intensities and selected areas of interest are quantified using functions “Straight”, “Rectangle”, “Line Width”, “Measure” in Fiji or “Measurement point”, “Surfaces” in Imaris. 3D representations are generated using “Surface visualization” in Imaris. Membrane volume quantifications are performed using an ImageJ script written by Christian Tischer at the Advanced Light Microscopy Facility at EMBL, which is described in detail and available at https://git.embl.de/grp-cba/glia-volume-measurement.

#### Selecting regions of interest (ROI)

In Fig. S4, ROI_1_ is defined from posterior end of 1st pharyngeal bulb until the anterior start of the second pharyngeal bulb, ROI_2_ from the anterior end of 1st pharyngeal bulb until the nose tip (based on bright-field images). In Fig. 5B-D and Fig.S5, ROI_1_ is from the posterior end of 1st pharyngeal bulb to anterior of 2nd pharyngeal bulb, ROI_2_ from the posterior end of 2nd pharyngeal bulb. In Fig. 5E, UNC-52 signal intensity is obtained on a line parallel to animal’s ventral side. In Fig. 6A-D, the distance between DLG-1 AJM-1 is measured between the centers of the 2 signals. In Fig. 6C-D, DLG-1 position is measured with 2nd pharyngeal bulb as external reference. In Fig. 6E-F, distance is from DLG-1 enrichment to the neighboring signal of UNC-52 accumulation.

#### For analysis of time-lapse imaging

Quantifications of glia length are performed using the function “Measure” in Fiji software. Videos are made using Imaris Animation feature. Images have been centered around the centroid of the posterior CEPsh glia in a max-intensity projection. Image alignment, registration, and convertion to movies using a Python code available upon reasonable request.

#### For imaging with Brillouin microscopy

Images are acquired as described above.

Spatial maps of elasticity and viscosity are plotted from the acquired data, and function “Measure” and Lookup table mpI-Inferno in Fiji and adjusting the “Brightfield and Contrast” at 7,5 - 8,2 or 1,02 - 2,05 respectively. ROI is selected with the function “Polygon selections” in Fiji. ROI excludes the pharynx to avoid signal variability. From the raw Brillouin shift and FWHM linewidth values we compute the Brillouin elastic and viscous contrast analogous to Antonacci et al. 2020.

### Statistical analysis

Sample sizes and statistical tests are chosen based on previous studies with similar methodologies; the data met the assumptions for each statistical test performed. No statistical method is used to decide sample sizes. All population experiments are performed at least three times using at least

100 individuals in total, comprised of biological replicates and yielding similar results. Independent transgenic lines or individual days of scoring for mutants are treated as independent experiments for the standard deviation. Chi-square test (GraphPad) is used to quantify statistical significance of all phenotypes in population studies, heat-shock experiments, RNAi experiments, drug applications, liquid culture experiments, contingency tables for defect correlations. The student’s t-test (GraphPad) is used for comparison of normally distributed values of glia size, nuclei distances, axon lengths, synaptic sites/contacts areas and intensities, signals of UNC-52, DLG-1, AJM-1, and Brillouin values. For all results, mean ± standard deviation (s.d.) is represented, unless otherwise noted. P values are calculated using GraphPad. The t ratio for a paired t-test presents the mean of these differences divided by the standard error of the differences. The number of degrees of freedom equals the number of pairs minus 1.

### Blinding and randomization during data analysis

Blinding during data analysis is not performed. Samples are allocated to groups of the genetic background (genotype), detected by standard genetic/genomic approaches. Samples are randomly selected within these groups, based on previous studies with similar methodologies. Phenotypic analysis is performed by more than one individual researcher for experiments of RNAi screens and image quantifications CEPsh glia sizes and age-dependent defects.

## Acknowledgments

We thank colleagues in the Developmental Biology Unit at EMBL Heidelberg, Michel Labouesse, Alba Diz-Munoz, Michael Dorrity, Lena Kutscher, Jean-Louis Bessereau, Shai Shaham Anne Ephrussi, Cornelia Bargmann,sfor scientific input, the Heiman and Shaham labs for sharing reagents. Some strains were provided by CGC, funded by NIH (P40 OD010440). Wormbase (www.wormbase.org) was used for experimental design and analysis. We thank the Advanced Light Microscopy Facility, and Christian Tischer, at EMBL, for invaluable technical help. R.P. acknowledges support of an ERC Consolidator Grant (no. 864027, Brillouin4Life), and the German Center for Lung Research (DZL). This work was supported by the European Molecular Biology Laboratory.

## Author Contributions

G.R. and F.C. conceived the project. F.C., M.B., F.C., S.R., and G.R. performed all experiments. G.R., C.B. and R.P. designed the Brillouin experiments. G.R, F.C. and C.B. performed Brillouin experiments. G.R. led the project and wrote the paper together with F.C. and input from all authors.

