## Supplemental Figures Legends for "An interplay of HSP-proteostasis, biomechanics and ECM-cell junctions ensures *C. elegans* astroglial architecture"

### Supplementary figures legends

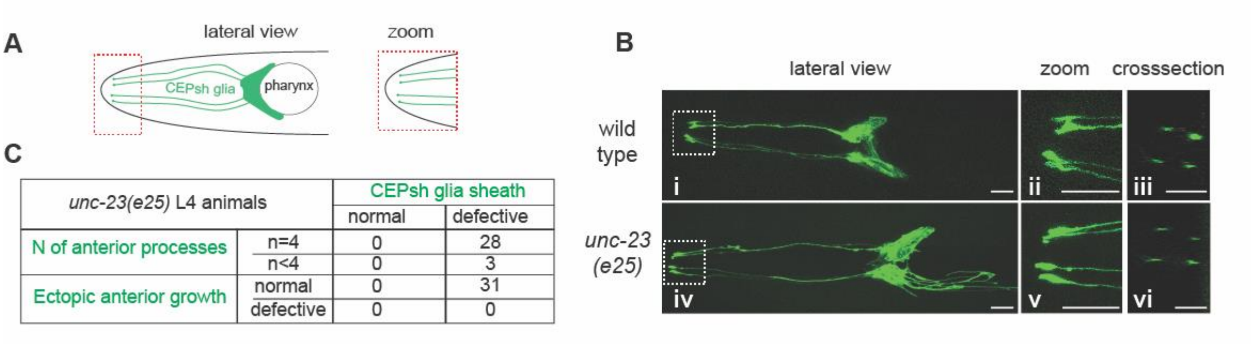


**Figure S1.**

**The anterior processes of CEPsh glia are not defective in *unc-23* mutant larvae.**

**A.** Each of the 4 CEPsh glia has an anterior process and tip arriving to C. elegans sensory endings. B. Anterior processes and tips of CEPsh glia (green) are largely not defective in *unc-23* L4 larvae mutants compared to wild-type animals, in zoom view (**i-vi**) and cross-sectional view (**iii-vi**). Scale bars, 10µm. Animal axes as in Figure 1. **C.** Quantification of CEPsh glia anterior processes number/ defects in *unc-23* mutant L4 animals, performed as in Methods.

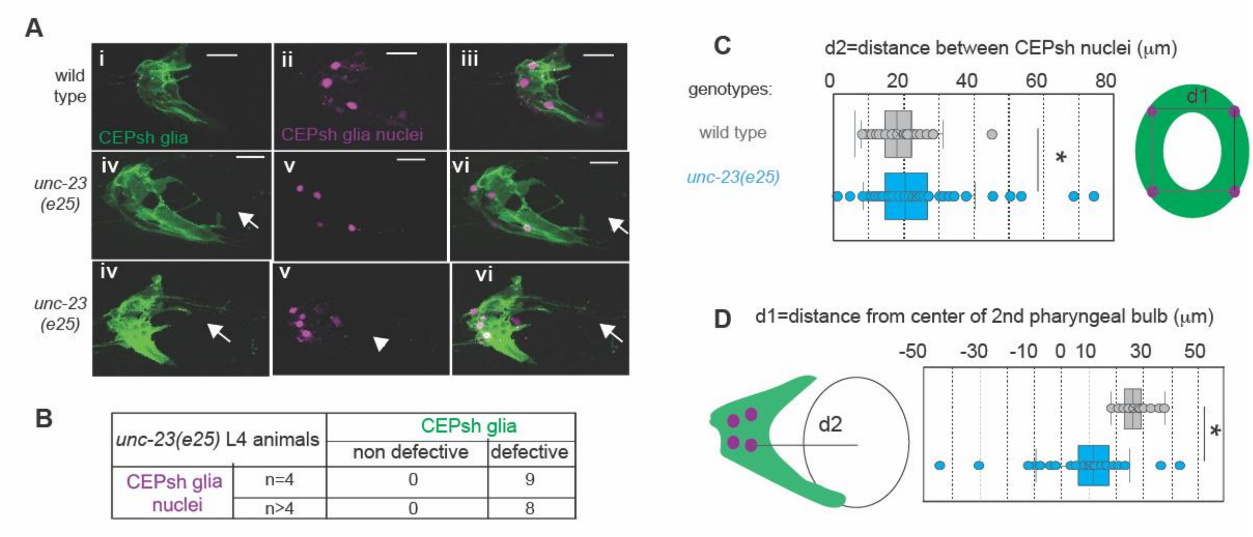


**Figure S2.**

***unc-23* mutant L4 animals present fragmentation and mispositioning of CEPsh glia nuclei (A-C).** Compared to wild-type animals L4 *unc-23* mutant animals with established defects in CEPsh glia membrane sheath (green) may present fragmentation of CEPsh glia nuclei (magenta) (A-B). CEPsh glia also present abnormal placement of nuclei between each other (C) and relative to the second pharyngeal bulb (D). This is in contrast to L2 *unc-23* mutant animals that present no defects in CEPsh glia nuclei (Figure 2). n=20 animals total per genotype. *, p-value<0,01, unpaired t-test.

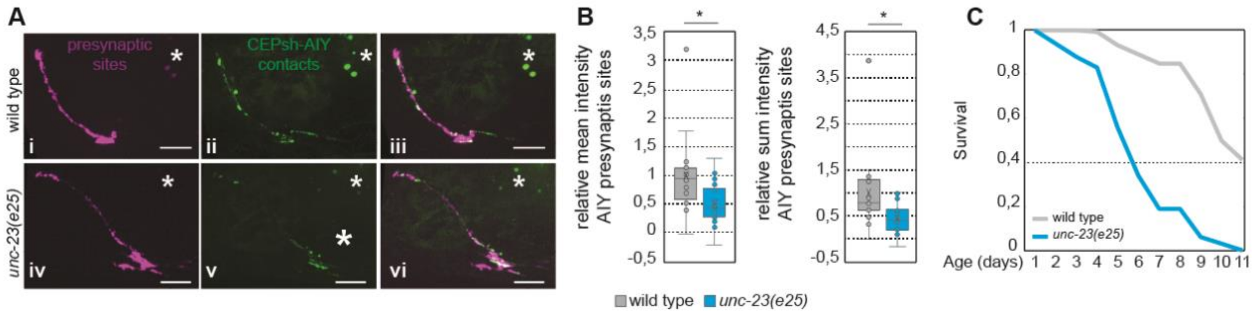


**Figure S3.**

**Integrity of presynaptic sites and aging are affected in *unc-23* mutant animals.**

**(A-C).** During aging, in five-day old adults, AIY presynaptic sites (magenta) have decreased mean and sum intensity in *unc-23* mutant animals (Ai-iii, **B**) compared to wild-type animals (Aiv-vi, **B**), * gut autofluorescence. n=12 animals per genotype. unpaired t-test. **C.** *unc-23* mutants show decreased longevity compared to wild-type animals. n=100 animals per genotype. Animal axes, P values as in Figure 3.

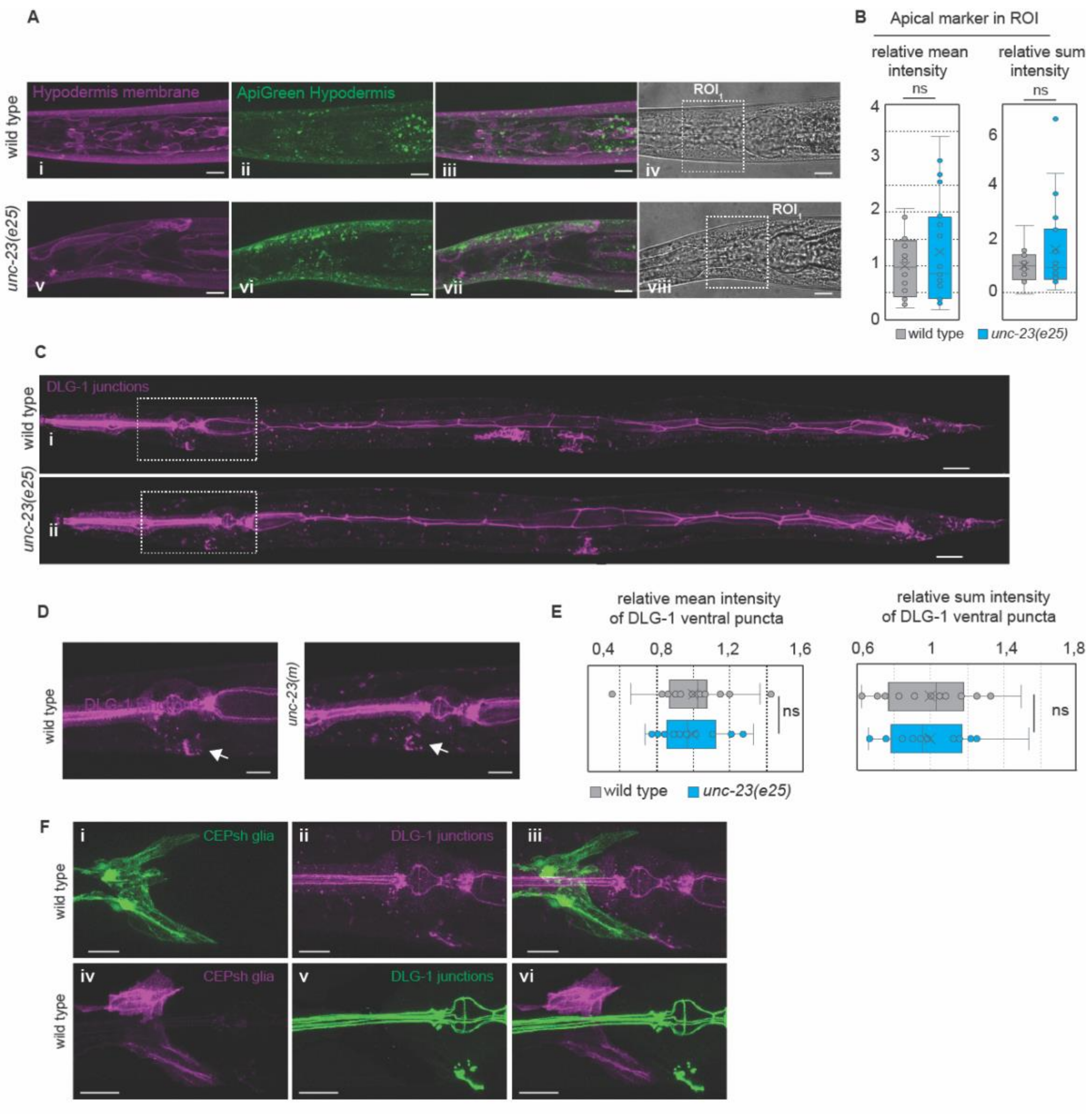


### Figure S4.

**UNC-23/BAG2 acts for glial polarity independently of epithelial polarity and integrity.**

Overall epithelial integrity is not disrupted in *unc-23* mutants, as assessed by epithelial membranes labeling across the body (magenta) (Ai,v,B), and quantitative analysis of the epithelial apical domain marker ApiGreen (green), neighboring the CEPsh glia (Aii,vi,B). **C-E.** Epithelial integrity of DLG-1 junctions (magenta) throughout the body is not disrupted in *unc-23* mutants (C) and areas of DLG-1 enrichments are comparable in *unc-23* mutants and wild-type animals (D,E). **F.** Epithelial DLG-1 junctions maintain close proximity to the posterior edge of the CEPsh glia membrane sheath in wild-type animals, as assessed by epithelial DLG-1 (Fi-iii) or endogenously-tagged DLG-1. The doted squares in A are regions of interest (ROI) in B, quantified as described in Methods. The doted squares in C are D panels. Molecular reporters used as listed in Methods and Tables S1, S3. n= 20 animals scored per genotype. *, p-value<0,01, unpaired t-test. ns, non-significant. Arrows, DLG-1 ventral enrichment. Scale bars, 10μm. Animal axes as in Figure 1.

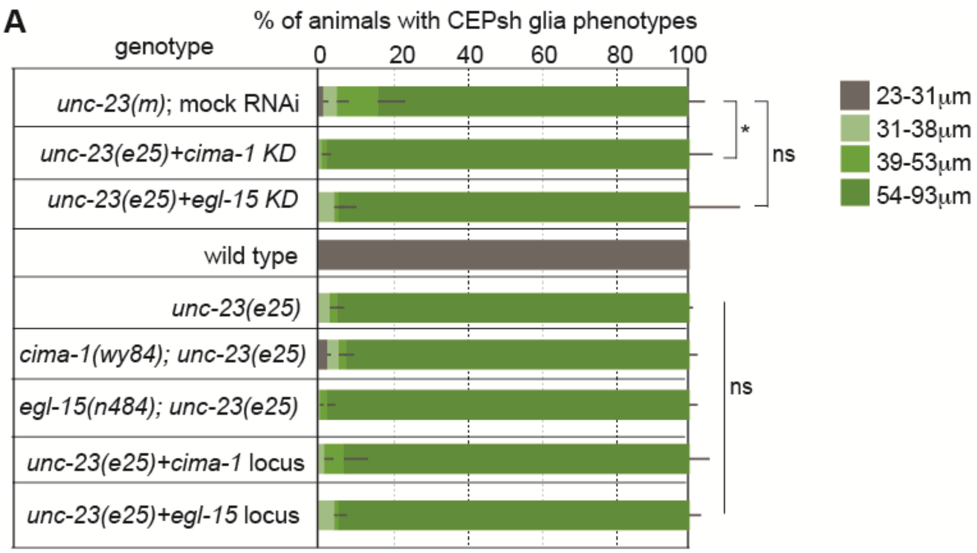


**Figure S5.**
**UNC-23/BAG2 acts for CEPsh glia integrity independently of epithelial allostery cues.**
**A.** RNAi-mediated knock-down (KD) of epithelial *allostery cues* EGL-15/FGFR and CIMA-1/ SLC17A5 (as defined by Fan et al, 2020), or their genetic mutants or their overexpression by formid expression (locus) do not suppress the CEPsh glia defects in *unc-23* mutants. * p value< 0,001 (RNAi of cima-1 appears to enhance the CEPsh glia defects in *unc-23* mutants). Error bars, mean ± standard deviation, n≥ 3 independent experiments, animals per genotype: n≥ 150 animals. Chi-square test.

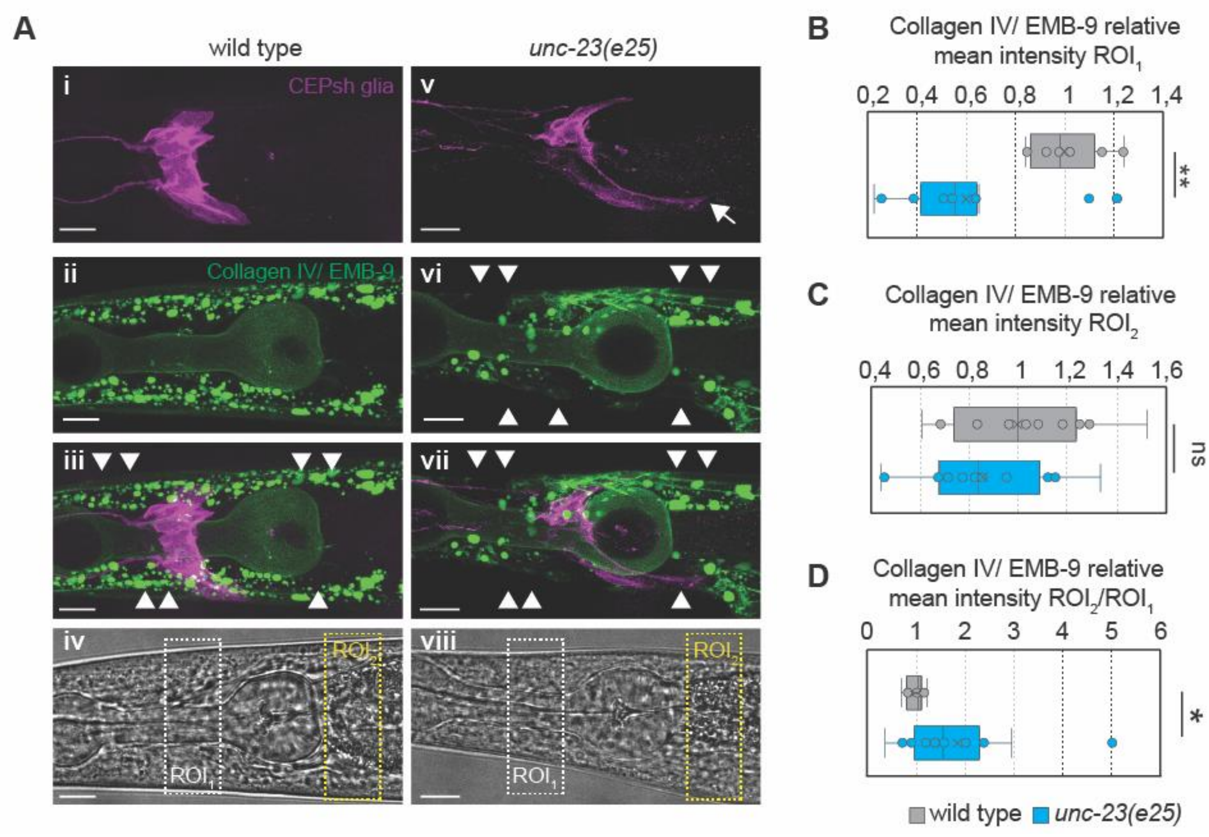


**Figure S6.**

**Collagen IV/ EMB-9 localization is impaired in *unc-23* mutants.**

**A-D.** EMB-9 (magenta) localization is impaired in *unc-23* mutants compared to wild-type animals, accumulating in the posterior part of CEPsh glia (green). Scale bars, 10μm. Arrows, CEPsh glia defects. Arrowhead, EMB-9 defects. $ROI_1$, $ROI_2$ correspond in A-D. Animal axes as in Figure 1. n=20 animals per genotype. *, p-value<0,05, unpaired t-test. ns, non-significant.

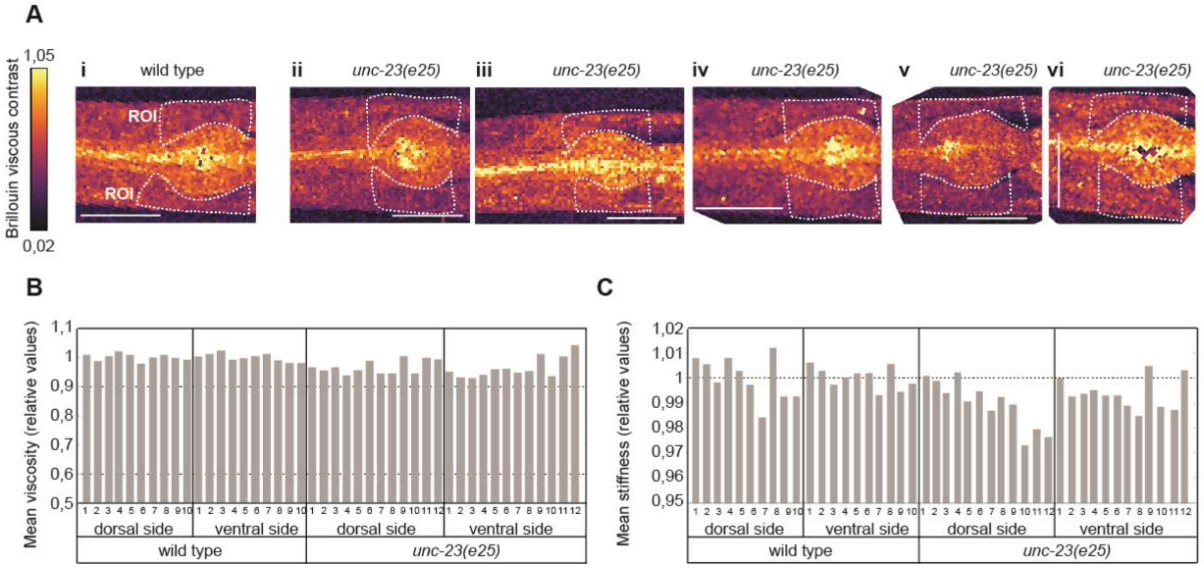


**Figure S7.**

**Brillouin measurement of viscosity and elasticity of wild-type and *unc-23* mutant individuals**

**A-B.** *unc-23* mutants show a decreased Brillouin viscous contrast of the tissues compared to wild-type animals, in regions neighboring CEPsh glia localization. Images (A) of individual wild-type and mutant animals, and measurements as values relative to the wild-type (A). **C.** *unc-23* mutant's tissues show a higher variability in the Brillouin elastic contrast compared to the wild-type in regions neighboring CEPsh glia localization. Scale bars, 10μm. White dotted rectangles in (A), ROI in (B,C). Values are depicted relative to the mean value of Brillouin contrasts in wild-type animals.
